# Data Heterogeneity Limits the Scaling Effect of Pretraining in Neural Data Transformers

**DOI:** 10.1101/2025.05.12.653551

**Authors:** Linxing Preston Jiang, Shirui Chen, Emmanuel Tanumihardja, Xiaochuang Han, Weijia Shi, Eric Shea-Brown, Rajesh P. N. Rao

## Abstract

A key challenge in analyzing neuroscience datasets is the profound variability they exhibit across sessions, animals, and data modalities–i.e., heterogeneity. Several recent studies have demonstrated performance gains from pretraining neural foundation models on multi-session datasets, seemingly overcoming this challenge. However, these studies typically lack finegrained data scaling analyses. It remains unclear how different sources of heterogeneity influence model performance as the amount of pretraining data increases, and whether all sessions contribute equally to downstream performance gains. In this work, we systematically investigate how data heterogeneity impacts the scaling behavior of neural data transformers (NDTs) in neural activity prediction. We found that explicit sources of heterogeneity, such as brain region mismatches among sessions, reduced scaling benefits of neuron-level and region-level activity prediction performances. For tasks that do exhibit consistent scaling, we identified implicit data heterogeneity arising from cross-session variability. Through our proposed session-selection procedure, models pretrained on as few as five selected sessions outperformed those pretrained on the entire dataset of 84 sessions. Our findings challenge the direct applicability of traditional scaling laws to neural data and suggest that prior reports of multi-session scaling benefits may need to be re-examined in the light of data heterogeneity. This work both highlights the importance of incremental data scaling analyses and suggests new avenues toward optimally selecting pretraining data when developing foundation models on large-scale neuroscience datasets.

## 1. Introduction

Recent advances in foundation models have revolutionized the modern machine learning paradigm. Across domains such as language and vision, it has been shown that “pretraining” a generic model on large-scale data before “finetuning” it to the actual tasks achieves much better performance than task-specific models (Devlin et al., 2019; Brown et al., 2020; Chung et al., 2024). This success has inspired similar efforts in systems neuroscience, where the goal is to develop foundation models trained on large, multi-session, multi-animal neural datasets of neural activity recordings. However, neural recordings pose unique challenges: data collected across brain regions, sessions, and individuals often exhibit substantial variability (Laboratory et al., 2021; 2025; Waschke et al., 2021). Even within the same recording session, stochasticity of neuronal firing and uncontrolled behavior can lead to significant trial-to-trial variability (Harris & Thiele, 2011; Stringer et al., 2019; Peterson et al., 2021). Furthermore, neural data can be non-stationary due to synaptic plasticity that induces gradual changes in population dynamics across days (Rule et al., 2019; Driscoll et al., 2022). These challenges raise a key question: Can neural foundation models overcome these sources of heterogeneity and learn more generalizable representations with more pretraining data?

While several recent studies have demonstrated performance gains from multi-session pretraining on a wide range of encoding and decoding tasks, they typically lack fine-grained scaling analyses on the benefits of gradually increasing pretraining data (Azabou et al., 2023; 2024; Zhang et al., 2024b; 2025). Most comparisons are limited to models trained on single sessions versus entire datasets with few increments in the middle, making it unclear how different sources of heterogeneity impact performance scaling. Moreover, it remains unknown whether all pretraining sessions contribute equally to downstream performance improvements (see Appendix A for related work). As pretraining scales to thousands of sessions and hours of data (Ye et al., 2025; Azabou et al., 2024), understanding the scaling behaviors of the model becomes increasingly critical.

In this work, we systematically investigate how data heterogeneity affects the scaling behavior of neural data transformers (NDTs) (Ye & Pandarinath, 2021; Zhang et al., 2024b; Ye et al., 2025) using two large-scale datasets released from the International Brain Laboratory (Laboratory et al., 2025; 2024). These datasets differ in the consistency of recorded brain regions across sessions, offering us an opportunity to study how different levels of brain region heterogeneity in pretraining affect scaling. We further examine the effects of implicit heterogeneity such as session-to-session variability. Through a proposed session-selection procedure, we identify the impact of each pretraining session on downstream performance improvements. Our main findings include:

- We found that greater region-wise heterogeneity in pretraining data led to reduced improvements of neuron- and region-level activity prediction performances.
- To identify implicit heterogeneity, a session-selection procedure based on single-session finetuning performances can effectively identify most beneficial single sessions for pretraining.
- Models trained with as few as five selected sessions outperformed those with randomly chosen sessions even when the full dataset was used, demonstrating the impact of session-to-session variability in performance scaling.

Together, these findings suggest that previous claims regarding the scaling benefits of pretraining without detailed incremental experiments may be premature, pointing to the need for rigorous scaling analyses in future work on neural foundation models to accurately assess the promise of large-scale pretraining.

## 2 Experimental Setup

Figure 1 summarizes the experimental setup used throughout our study, which mostly follows Zhang et al. (2024b) whose experiments were conducted on a subset of the same RepeatedSite dataset we used. We discuss the datasets, training pipeline, and evaluation metrics in detail below. More details are included in Appendix B.

**Figure 1:**
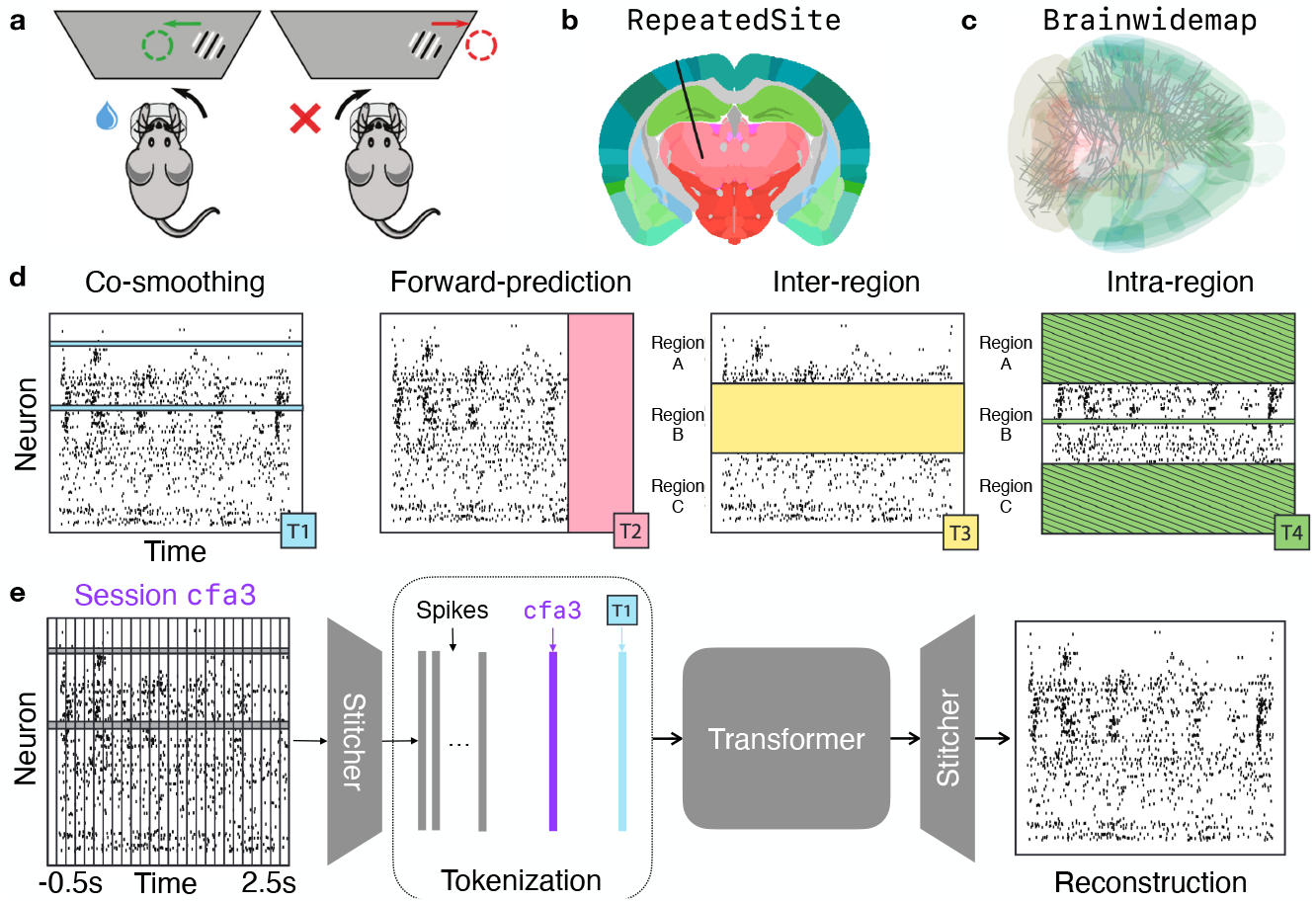
Experimental Setup. **(a)** Schematic of the visual decision-making task performed by mice. **(b)** Planned probe insertion location (black line) for all sessions in the RepeatedSite dataset. **(c)** Different probe insertion locations (gray lines) for different sessions in the Brainwidemap dataset. **(d)** Four different masking schemes of raw spike counts for model training. **(e)** The model architecture. Sub-figures adapted from Laboratory et al. (2021; 2025; 2024); Zhang et al. (2024b). See text for details.

### 2.1 Datasets

We used two multi-brain-region, multi-animal/session datasets from the International Brain Lab (IBL) collected from mice. Animals performed a visual decision-making task where they detected the presence of a visual grating (of varying contrast) to their left or right and rotated a wheel to bring the stimulus to the center (Fig. 1(a)). The main difference between the two IBL datasets lies in how often brain regions were repeatedly recorded across sessions. In the RepeatedSite dataset (henceforth RS), each session attempted to record from the *same* brain regions (Fig. 1(b), black line shows planned electrode insertion position). In contrast, the Brainwidemap dataset (henceforth BWM) aimed to cover as many *different* brain regions as possible (Fig. 1(c), gray lines show planned insertion positions), leading to little repetition of regions across sessions.

We used 89 out of 91 sessions in RS, excluding two sessions with fewer than one hundred trials. Five out of 89 sessions were held out for finetuning and evaluation. We randomly selected 200 sessions out of 460 sessions in BWM for pretraining and 10 sessions for finetuning and evaluation. Trials within each session were randomly split into training, validation, and test sets using an 8:1:1 ratio. Each trial included three seconds of neural activity, spanning from 0.5 seconds before to 2.5 seconds after stimulus onset with 20 ms bins for spike counts. The data from each session is thus a three-dimensional (trials × timesteps × neurons) tensor of integer spike counts.

### 2.2 Training pipeline

**Multi-masking scheme** During training, input spike count vectors were masked in one of four ways, as illustrated in Figure 1(d): (1) Co-smoothing: selected neurons’ activities are masked; (2) Forward-prediction: all neurons’ selected timesteps are masked; (3) Interregion: all neurons’ activities in a selected brain region are masked; (4) Intra-region: selected neurons in a selected brain region, along with all other neurons in other brain regions, are masked. Models were trained to reconstruct masked input from the unmasked (Devlin et al., 2019) by maximizing the log likelihood of the Poisson distribution, with the model outputs as the predicted firing rates. We also applied causal attention in NDT if inputs were masked with forward-prediction to ensure no future neural activities were used to predict the present. Zhang et al. (2024b) showed that this multi-masking scheme significantly outperforms a scheme using forward-prediction masks alone in spike prediction tasks.

**Model architecture** We used the neural data transformer (NDT) architecture by Ye & Pandarinath (2021) that has been widely applied to neural encoding and decoding tasks (Le & Shlizerman, 2022; Ye et al., 2025; Zhang et al., 2025). NDT also achieves state-of-the-art performance on the IBL dataset we used (Zhang et al., 2024b; 2025). Since different sessions have different numbers of neurons recorded, a session-specific linear layer (encoding “stitcher”) maps raw spike counts to spike embeddings (Fig. 1(e) left) whose dimensions are shared across sessions (Pandarinath et al., 2018). A session embedding and a masking scheme embedding are also appended to input sequences. Lastly, another session-specific linear layer (decoding stitcher) maps the output of the transformer back to reconstructed spike rates (Fig. 1(e) right).

### 2.3 Evaluation

**Baseline** To show the effect of scaling up pretraining data, we directly trained single-session models on the training set of each heldout session as the baseline models.

**Metrics** The bits-per-spike metric (BPS) is widely used to evaluate neural activity prediction performance (Rieke et al., 1999; Pei et al., 2021; Zhang et al., 2024b; 2025):

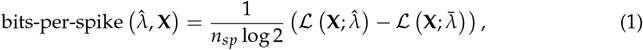

where 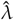 is the predicted spike rates by the model, **X** is the true spike counts, *n*_*sp*_ is the total spike count of **X**, ℒ is the log likelihood function of Poisson, and 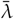 is the mean firing rate of **X**. The BPS metric essentially evaluates the goodness-of-fit statistics of a model over the null model, normalized by the spike counts. Changes in BPS directly reflect changes in model log likelihood 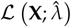 when evaluated on the same dataset, as other terms remain constant. For all experiments, we report the metrics on the test sets of the heldout sessions after finetuning models to their training sets.

**Tasks** Each masking scheme used for training corresponds to a leave-one-out evaluation task of activity prediction, namely: (1) Co-smoothing: Activities of each neuron were predicted from all other neurons; (2) Forward-prediction: The model predicted neural activities in continuous, non-overlapping windows of 200 ms at a time (10 timesteps) given previous ground truth activities, starting from stimulus onset to 2.2 seconds after stimulus onset (110 timesteps in total); (3) Inter-region: Activities of all neurons in each region were predicted from all other regions; (4) Intra-region: Activities of each neuron in a region were predicted from all other neurons in that region. Repeat for each region.

## 3 Data heterogeneity limits NDT models’ scaling behavior

As mentioned, the difference in brain region overlaps between RS and `BWM` provides a natural setting to study how pretraining data heterogeneity affects model scaling behavior. `BWM` sessions contain neural activity from largely non-overlapping brain regions, resulting in greater single-neuron and brain-region heterogeneity compared to RS. We hypothesize that increased heterogeneity in `BWM` will reduce the scaling benefits from pretraining on more sessions.

We conducted our scaling analysis as follows. Using the RS dataset, we pretrained NDT models on 20, 40, 60, and 84 sessions, then finetuned them to each of five heldout sessions. For `BWM`, we pretrained models on 20, 40, 60, 80, 100, 150 and 200 sessions (out of 460 total), then finetuned them on ten heldout sessions. During finetuning, a new session embedding and two session-specific stitchers were learned from scratch while the mask embedding and the core NDT parameters were initialized from pretraining.

### 3.1 Performance scaling is weaker in BWM than RS

Before analyzing whether pretrained models’ performances scale with more pretraining data, we first confirm they indeed outperformed baseline models (Appendix C). Next, we investigate whether the task performances scale with increased pretraining data. Figure 2(a) and (b) show the evaluation results of models trained on RS and `BWM` data, respectively. Although performance generally improved with more pretraining data, the scaling effects were relatively modest. Figure 2(c) illustrates the percentage performance gains relative to the 20-session model, with the largest improvement of 4.2% observed on the intra-region task on RS using more than four times the amount of data. Notably, performance gains reduced across the co-smoothing, inter-region, and intra-region tasks on `BWM` compared to RS. For co-smoothing in particular, scaling benefits were negligible in `BWM`. In contrast, the forward-prediction performance scaled consistently on the two datasets. However, the forward-prediction performance also plateaued around 80 pretraining sessions, suggesting a potential upper limit on the performance improvements achievable through pretraining given the current scope of data. These results support our hypothesis that greater brain region heterogeneity in `BWM` limits the effectiveness of pretraining, particularly on single-neuron and region-level tasks.

**Figure 2:**
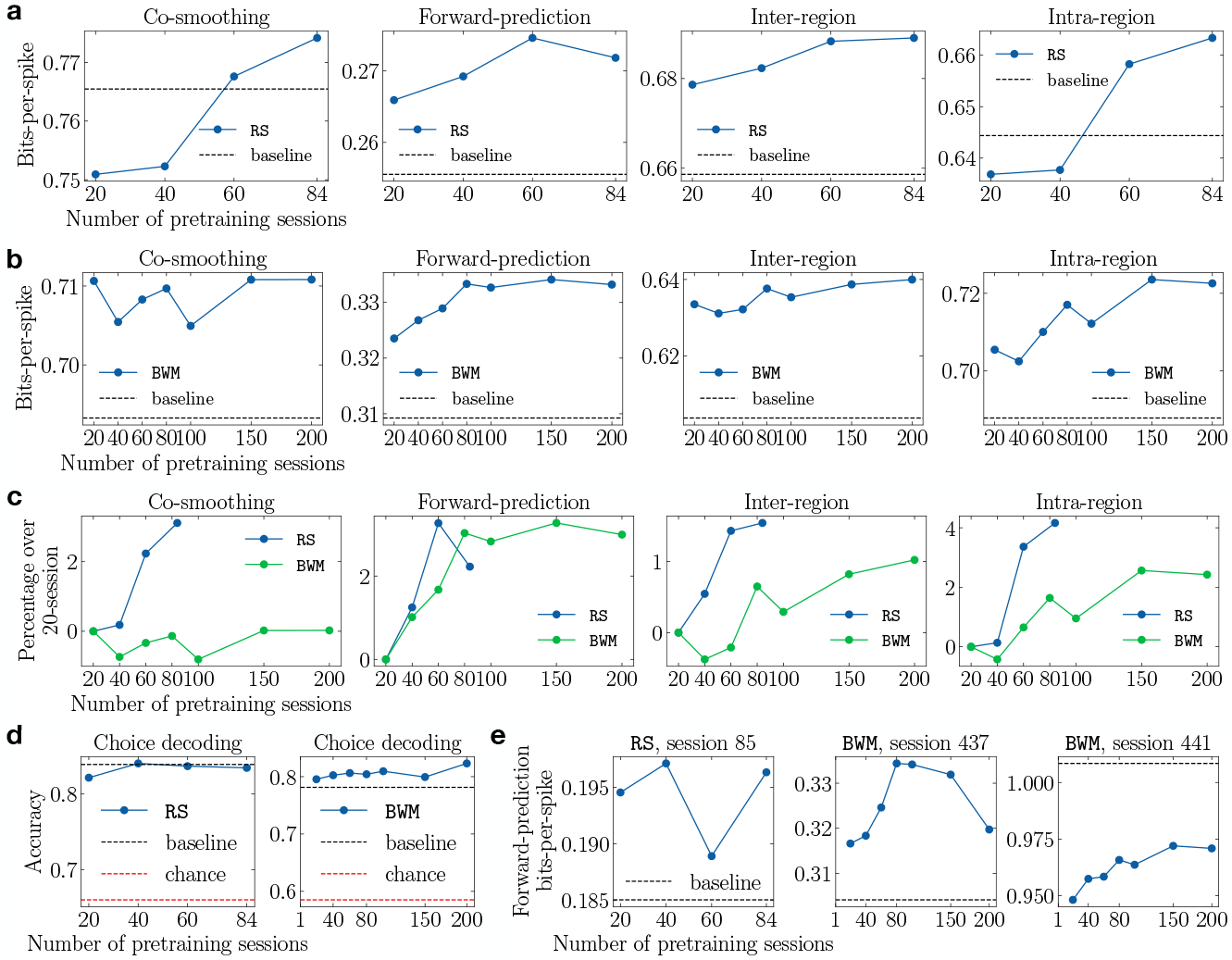
Scaling Behavior of NDT Models. Plots show pretrained NDT models’ finetuning performances as the number of pretraining sessions increases. **(a)** Performances of each neural activity prediction task on RS data. Black dashed lines show the baseline models’ performances. **(b)** Same as (a) but on `BWM` data. **(c)** Percentage improvements of models pretrained with more sessions over the 20-session model. **(d)** Choice decoding performances on RS (left) and `BWM` (right). Red dashed line shows the chance prediction accuracy. **(e)** Forward-prediction performance examples on individual heldout sessions from RS (first panel) and `BWM` (second & third panels).

To probe the quality of the NDT’s internal representations, we trained a logistic regression classifier on the output of the third intermediate transformer block (out of five, see Appendix B) to predict the animals’ decisions. Previous work typically performed such decoding analyses using reconstructed spikes rather than models’ internal representations (Pei et al., 2021; Zhang et al., 2024b). Under the latter setup, improvements in choice decoding performance could reflect better spike reconstruction rather than better representations of choice-related latent states underlying the data. With our setup, decoding accuracies are higher in RS than `BWM` (Appendix C). Figure 2(d) shows the changes in classification accuracy with increasing amounts of pretraining data. There was no clear scaling of decoding performance on either dataset. Taken together, these results highlight the importance of investigating the scaling behaviors of pretrained models with more fine-grained data increments.

We also observed large cross-session variabilities in the finetuning performances. Fig. 2(e) shows the forward-prediction performance of three heldout sessions (see Appendix D for all sessions). In addition to the substantial differences in absolute bits-per-spike values, their scaling trends deviate from the session averages (second panel in Fig. 2(a) & (b)). Such variability indicates a more implicit form of data heterogeneity that comes from individual differences among animals. Given this observation and our previous findings on the pretrained models’ limited scaling behavior, a natural question arises: **Can we identify more beneficial sessions than others in the pretraining dataset for improving scaling performance?** We answer this question in the next section.

## 4 Identifying more beneficial single sessions for performance scaling

We hypothesize that each pretraining session exhibits varying degrees of distribution shift relative to a heldout session. We call this implicit data heterogeneity, which arises from subtle individual differences among animals and sessions that are harder to identify than explicit sources of heterogeneity such as task design and brain regions. We expect that models pretrained on sessions “closer” to the heldout sessions will achieve higher performances more data-efficiently than models pretrained with randomly selected sessions.

To test this, we conducted our experiments on the RS dataset, which allows us to control the brain region heterogeneity as discussed in the last section. We trained NDT models with only forward-prediction masking to focus on this evaluation task, which exhibits the most consistent scaling behavior on the two datasets (Fig. 2(c)). For more fine-grained scaling analysis, we pretrained models on 1, 2, 3, 4, 5, 10, 20, 30, 40, and all 84 sessions.

### 4.1 Ranking pretraining sessions by single-session finetuning performances

First, we propose using single-session finetuning performances as an estimate of the “closeness” between the data distributions of a pretraining session and a heldout session. Figure 3 illustrates this process: during the pretraining stage, we trained 84 single-session models, one for each pretraining session (Fig. 3(a)). During the finetuning stage, for a particular heldout session, we trained two new stitchers (for encoding and decoding) for each of the pretrained transformers while keeping the transformers’ weights frozen (Fig. 3(b)). This ensures the finetuning performance maximally depends on the features learned from the pretraining session, as the only adjustable weights were the input/output linear layers that map the raw spike counts to the frozen feature space and back. Lastly, we report each model’s forward-prediction performance on the heldout session’s validation set, yielding 84 metric values – one per pretraining session. The pretraining stage (Fig. 3(a)) was performed once, while the finetuning and ranking stage (Fig. 3(b)) were repeated for each heldout session. See Appendix E for the ranked single-session finetuning performances.

**Figure 3:**
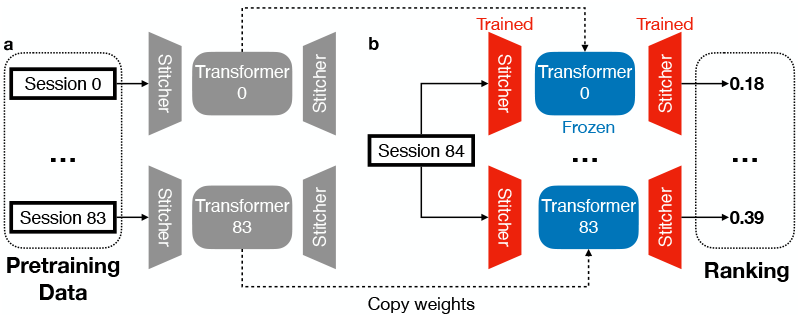
Schematic of the ranking process. **(a)** Pretraining stage: We trained 84 single-session models, each consisting of a transformer and two sessionspecific stitchers. **(b)** Finetuning stage: For each pretrained model, we trained two new stitchers on the heldout session’s training set, keeping the transformer weights frozen. Models were ranked by their bits-perspike metric on the heldout session’s validation set.

We conducted our data scaling experiments by incrementally selecting more pretraining sessions in three session-selection orders: random, ranked (based on the procedure above), and reverse-ranked. To reduce variance, we used three random seeds for both ranking sessions and training models in the scaling analysis, including the baseline models that were directly trained on the heldout sessions. For the random order, we used three different shuffles of the session list. The transformers’ weights were frozen for all finetuning experiments across data orders to be consistent with the ranking procedure. We limited experiments to a maximum of 40 pretraining sessions (except for the full 84-session case) since more selected sessions overlap as we exhaust the pretraining data.

### 4.2 Pretraining on five top-ranked sessions outperforms all random sessions

Figure 4(a) shows the performances of our scaling analysis on each heldout session’s test set with different session orders (see Appendix G for qualitative examples). The results clearly show that in all heldout sessions, models pretrained with ranked session order outperform those trained with randomly chosen sessions. Importantly, the models pretrained with reverse-ranked sessions achieved worse performances than random-order models, proving the validity of our ranking procedure based on single-session finetuning. Notably, the performance differences in ranked, random, and reverse-ranked settings are more pronounced in low-data regimes (fewer than ten sessions). Since the number of trials was different among sessions, we also plotted the model performances in Figure 4(a) against the total number of trials from the pretraining sessions for fairer comparisons. As shown in Figure 4(b), the same performance differences hold among the different session-selection orders given the same number of trials. We fit the models’ performances using linear regression (with logarithmic input). In contrast to the success of “scaling laws” in machine learning (Kaplan et al., 2020; Hoffmann et al., 2022; Muennighoff et al., 2023), the actual pretraining performance using the entire 84 RS sessions (Fig. 4(b) red stars) is consistently lower than the extrapolated performance (Fig. 4(b) dashed lines), indicating limited scaling effects for the neural IBL data with the NDT model. This further supports our hypothesis that differences in pretraining and finetuning data distributions greatly affect the promises of neural data scaling.

**Figure 4:**
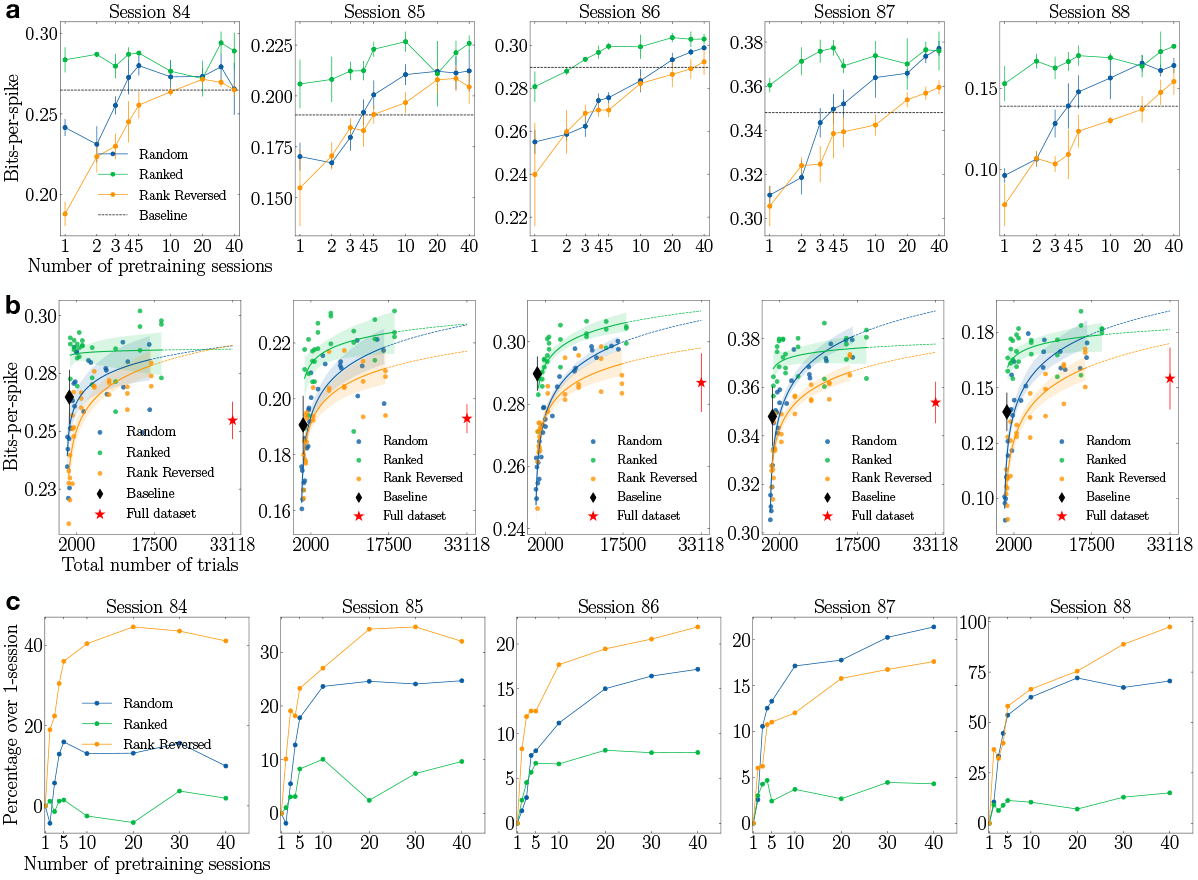
Scaling performances under different session orders. **(a)** Forward-prediction performances of each heldout session as we increased pretraining sessions according to random (blue), ranked (green), or reverse-ranked (orange) order. The error bars show the standard deviation over three seeds (ranked/reverse-ranked) or three shuffled orders (random). Black dashed lines show the baseline models’ performance (averaged over three seeds). **(b)** Same as (a) but with the total number of trials as the *x*-axis. Linear regressions were fitted with logarithmic *x* values and dashed lines show extrapolated predictions. Shading shows the standard deviations. Red stars show the performances of the models pretrained with all pretraining sessions. **(c)** Percentage improvements of models pretrained with more sessions over the 1-session model.

Table 1 summarizes the best percentage improvements over the baseline models for each session selection order, along with the performance of models pretrained on five top-ranked sessions. On average, models using rank-ordered session data achieved a 4% greater improvement over the baseline than models using random-ordered session data. Remarkably, models trained on just five ranked sessions outperformed the best models trained on randomly selected sessions, indicating an over 8× gain in data efficiency (compared to 40 random session models, which outperformed the models trained on all sessions (Fig. 4(b)). However, this also implies a reduced scaling effect compared to randomly selected sessions. Figure 4(c) demonstrates that the percentage performance gains using more pretraining data relative to using one pretraining session under each session selection order. The scaling effects when using the ranked sessions were clearly weaker than when using random or reverse-ranked sessions. Indeed, models with five ranked sessions already achieved 86% of the best model performances with all 40 ranked sessions (Table 1), suggesting that most of the pretraining benefit is concentrated in the top few sessions. The top sessions in different rankings were also sufficiently different based on the finetuning session used (Appendix F). Taken together, our analysis suggests that apparent scaling benefits in multi-session datasets can be highly sensitive to the specific sessions selected, due to substantial individual differences across sessions. Thus, it is extremely important for studies that claim scaling benefits to show detailed experimental results with fine-grained data increments.

**Table 1:** Percentage improvements over baseline with different session selection procedures.

| Heldout session | Session selection order (best of all experiments) |  |  | Ranked (top 5 sessions) |
| --- | --- | --- | --- | --- |
|  | Random | Ranked | Reverse-ranked |  |
| 84 | 5.74% | **11.08%** | 2.54% | 8.69% |
| 85 | 11.33% | **18.87%** | 9.39% | 16.92% |
| 86 | 3.13% | **4.78%** | 0.89% | 3.39% |
| 87 | 8.37% | **8.43%** | 3.30% | 8.43% |
| 88 | 19.12% | **26.60%** | 10.95% | 22.44% |
| Average | 9.54% | **13.95%** | 5.42% | 11.97% |

## 5 Conclusion

Our results show that data heterogeneity in multi-session electrophysiology datasets fundamentally limits performance improvements expected from increasing pretraining data. Future work can focus on (1) scaling experiments with other architectures beyond NDT and different learning objectives, including supervised approaches; (2) different modalities of neural data such as calcium imaging and local field potentials ^1^, and (3) more computationally efficient session selection strategies. In conclusion, our results show that pretraining neural encoding models with more sessions does not naturally lead to improved downstream performance. We strongly advocate for rigorous scaling analyses in future work on neuroscience foundation models to account for data heterogeneity effects.

## Acknowledgement

We thank Shuchen Wu, Nick Steinmetz, Matt Golub, and Luke Zettlemoyer for discussions. This work was supported by National Science Foundation EFRI grant 2223495 (RPNR), a UW + Amazon Science Hub grant (RPNR), a Frameworks grant from the Templeton World Charity Foundation (RPNR) and the Air Force Office of Scientific Research under award number FA9550-24-1-0313 (RPNR). Any opinions, findings, and conclusions or recommendations expressed in this material are those of the authors and do not necessarily reflect the views of the funders. We gratefully acknowledge InVirtualis for their support and for providing the computational resources to this research.

## A Related Work

### A.1 Foundation models in neuroscience

Foundation models represent a paradigm shift in artificial intelligence, allowing large-scale models pretrained on internet-scale data to be efficiently adapted to various downstream tasks through finetuning. These models demonstrate remarkable capability to learn versatile representations via self-supervised objectives and adapt effectively to downstream tasks (Radford et al., 2018; 2019; Wei et al., 2021; OpenAI et al., 2024; Radford et al., 2021; Dosovitskiy et al., 2020; Kirillov et al., 2023; Reed et al., 2022). Motivated by these advances, the neuroscience community has begun adopting similar approaches, starting with noninvasive human neural data across modalities (Cui et al., 2024; Thomas et al., 2022; labs at Reality Labs et al., 2024). More recent work trained large attention-based models on invasive rodent and nonhuman primate data, such as POYO (Azabou et al., 2023), POYO+ (Azabou et al., 2024), and NDT model series (Ye & Pandarinath, 2021; Ye et al., 2023; 2025). Using calcium imaging data, Wang et al. (2025) explored combining recurrent architectures with attention modules for predicting neural activities from visual stimuli and locomotion.

More specifically, NDT models scaled attention-based models to multi-session spiking data by incorporating context embeddings and learning a shared latent space across sessions, enabling transfer to new recording conditions (Ye et al., 2023; 2025). Azabou et al. (2023) extended the multi-subject pretraining paradigm to primate data and proposed single-spike tokenization through a PerceiverIO architecture (Jaegle et al., 2021). Zhang et al. (2024b) employed multiple spike masking schemes on the IBL RS dataset, upon which we based our work. Similarly, Zhang et al. (2025) proposed a novel multimodal training and masking method, demonstrating improved performance from multi-session training over single-session models.

### A.2 Scaling behavior in foundation models for neuroscience

Foundation models in NLP have been empirically shown to follow scaling laws, where performance improves predictably with more data and parameters (Kaplan et al., 2020; Hoffmann et al., 2022). In neuroscience, similar scaling effects have been explored through many pretraining studies. Azabou et al. (2023) used primate data from motor and premotor cortices and demonstrated that pretraining on over 100 hours of data enables rapid adaptation to unseen sessions. Azabou et al. (2024) extended their model to rodent visual cortex data and presented benefits of pretraining on over a thousand sessions. Using the IBL dataset, Zhang et al. (2024a) showed that reduced-rank regression models trained on hundreds of sessions across diverse brain regions outperform session-specific models, indicating benefits from multi-session data. Zhang et al. (2025) similarly reported performance gains from their multi-task masking strategies when pretrained across multiple sessions. While these results suggest that pretraining on multi-session data is generally beneficial, they often lack fine-grained analyses of how performance trends evolve with incremental pretraining data. As shown by our results in Fig. 4, scaling behaviors of the model may drastically vary when pretrained on different subsets of the pretraining dataset. Therefore, it may be misleading to conclude that the model enjoys scaling benefits with just a few data increments. In fact, recent studies have begun to observe limited scaling effects in foundation models for motor decoding when finetuning data exceeds 100 minutes (Ye et al., 2025), similar to our results.

## B Experimental setup details

In this section, we detail our experimental setup introduced in Section 2.

### B.1 Model architecture

Our model follows the architecture of Zhang et al. (2024b). At each time step *t*, a raw spike count vector 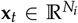 from session *i* (with *N*_*i*_ neurons recorded) is projected via a session-specific linear layer to a spike token with dimension *d*. Another linear layer with Softsign activation maps the spike tokens to embeddings. A session embedding and a masking scheme embedding were appended to the spike embedding sequence. Learned position embeddings are added to the input embeddings, making up the final input to the transformer block. Lastly, another session-specific linear layer maps the transformer output back to the spike count vectors of dimension *N*_*i*_ for session *i*.

### B.2 Hyperparameter selection

We tuned learning rates, dropout rates, weight decay rates, and batch sizes using small-scale experiments on single- and ten-session models. Optimal values were chosen based on test losses from pretraining sessions in RS. Other hyperparameter values were inherited from the implementation of Zhang et al. (2024b). Table 2 summarizes the hyperparameters we used for all experiments. We increased the spike token dimension to 1024 for `BWM` experiments to account for higher numbers of neurons. We used seed 42 for all experiments in Section 3, and seeds 10, 20, and 42 for experiments in Section 4 that required three seeds.

**Table 2:** Hyperparameter values across experiments.

| Hyperparameter | Value |
| --- | --- |
|  | Model |
| Spike token dim | 668 |
| Embedding dim | 512 |
| Feedforward dim | 1024 |
| # attention heads | 8 |
| # transformer blocks | 5 |
| Dropout rate | 0.2 |
|  | Training |
| Optimizer | AdamW Loshchilov & Hutter (2018) |
| Learning rate | 1e-4 |
| Learning rate scheduler | OneCycle Smith & Topin (2018) |
| Weight decay | 0.01 |
| Batch size | 16 |
| Gradient clipping | 1 |
| # Epochs | 1000 |
| Masking ratio | 0.3 |

The two session-specific linear layers contain approximately 1.2 million parameters on average per session. The number of parameters in NDT (shared across sessions) is roughly 12 million with the values in Table 2, which achieved better performance than smaller 8 million models and larger 24/38 million models on RS. We did not run larger models on `BWM` due to computing resource constraints.

### B.3 Compute

All models were trained on a single Nvidia A40 or L40 GPU. Single-session training and finetuning take about one hour to train on average. The full 84-session model on RS takes about 3.5 days, while the full 200-session model on `BWM` takes about a week to train. Finetuning jobs in Section 4 are significantly faster since we only train the two linear stitchers, with each taking about 20 minutes to finish on average.

## C Pretraining offers performance improvements across tasks over baseline models

Table 3 presents the evaluation metrics achieved by the baseline models and the best pretrained-then-finetuned models on RS and `BWM`, averaged over all heldout sessions. Pretraining improves performance on both datasets, though gains are smaller for co-smoothing and choice decoding tasks than others. Notably, co-smoothing metrics are higher than intra-region in RS but not in `BWM`, suggesting that models trained on RS data benefited more from cross-region information. This is consistent with the distinct recording strategies of the two datasets.

**Table 3:** Evaluation metrics of baseline and best pretrained models on RS and BWM. Higher values indicate better performance.

| Task | RS |  | BWM |  |
| --- | --- | --- | --- | --- |
|  | Baseline | Pretrained | Baseline | Pretrained |
| Co-smoothing (BPS) | 0.765 | **0.774** | 0.693 | **0.711** |
| Forward-prediction (BPS) | 0.256 | **0.275** | 0.309 | **0.334** |
| Inter-region (BPS) | 0.659 | **0.689** | 0.604 | **0.640** |
| Intra-region (BPS) | 0.644 | **0.663** | 0.688 | **0.724** |
| Choice decoding (accuracy) | 0.839 | **0.841** | 0.781 | **0.823** |

## D Cross-session variabilities in scaling behaviors

Here, we show detailed experimental results of the scaling analysis in Section 3. Figure 5 and Figure 6 show the finetuning performances on each heldout session in RS and `BWM`, respectively. As discussed in Section 3, there exist large cross-session variabilities in the finetuning performances among heldout sessions. These results suggest differential contributions of different sessions in pretraining, which we investigated in Section 4.

**Figure 5:**
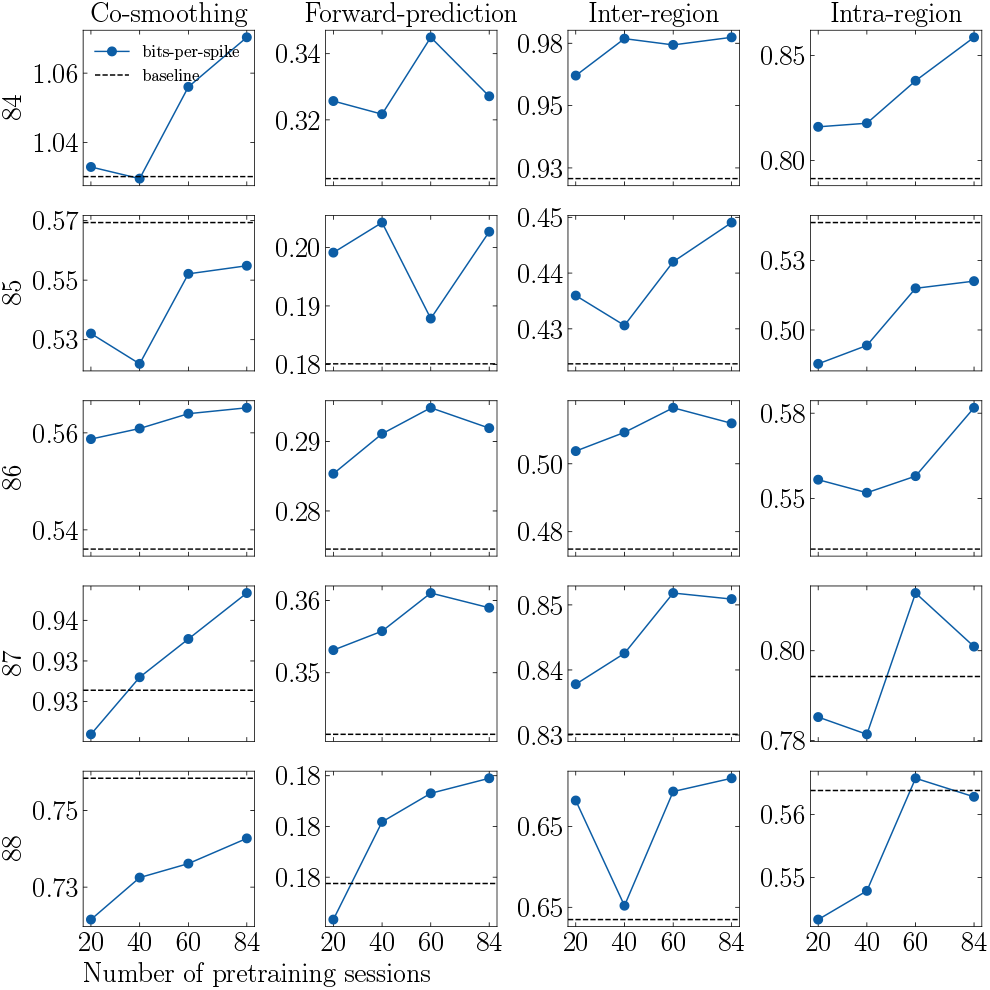
Pretrained models’ finetuning performances on each heldout session in RS. Black dashed lines show the baseline models’ performances.

**Figure 6:**
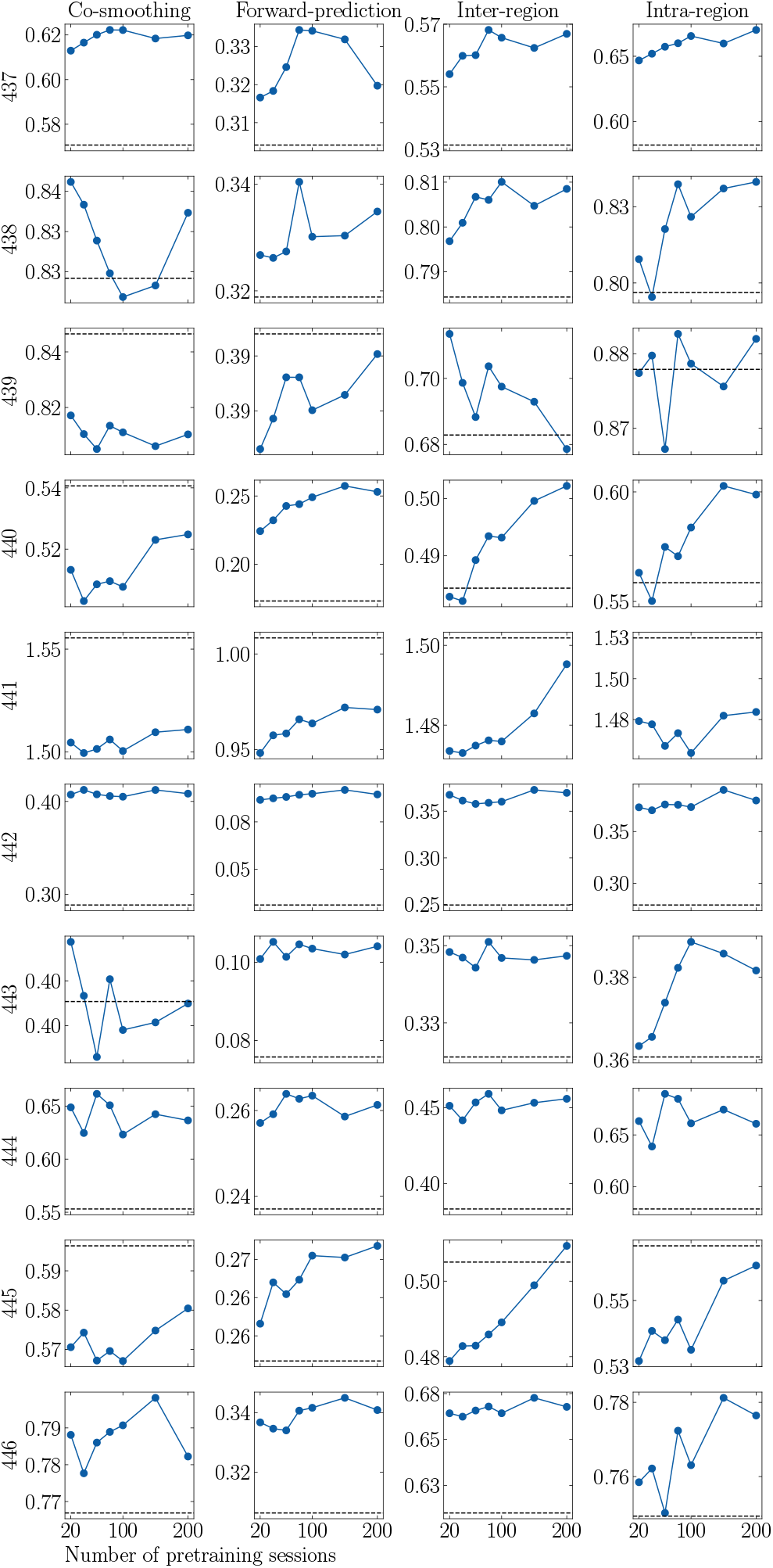
Pretrained models’ finetuning performances on each heldout session in BWM. Black dashed lines show the baseline models’ performances.

## E Finetuning results from the session-selection procedure

Here, we show the results of the session-selection procedure described in Section 4.1. Figure 7 shows the single-session finetuning performances on the validation set of each heldout session, sorted from high to low. As the figure shows, there exist large differences between the best and worst single-session finetuning performances, which are more noticeable in Sessions 84, 85, and 88 than in others.

**Figure 7:**
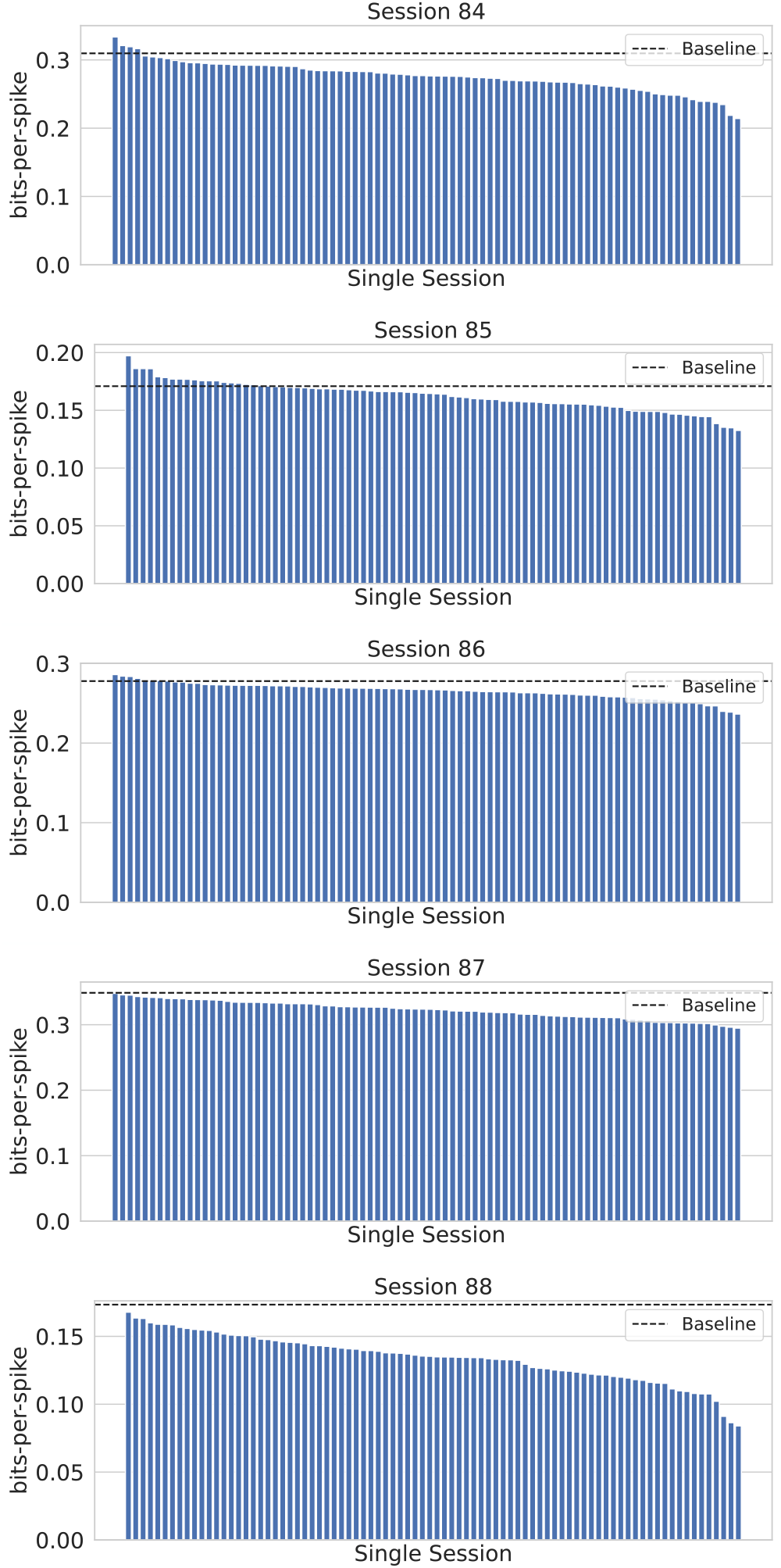
Single-session finetuning performances on the validation set of the heldout sessions from the session-selection procedure. Black dashed lines show the baseline models’ performances.

## F Session-specificity of the rankings

Here, we examine how session-specific the rankings were. Figure 8 shows the number of sessions shared in all five top-*k* ranking of the heldout sessions. The top 20 sessions were highly session-specific: no session appeared in the top five for all held-out sessions, and only three were shared in the top 23. In contrast, rankings became increasingly similar beyond 23 sessions, with 43 sessions shared in all five top 53 rankings. These results suggest that only a small number of top-ranked sessions have a strong impact on performance, while the remaining pretraining sessions are more consistent across held-out sessions and affect model performance less significantly.

**Figure 8:**
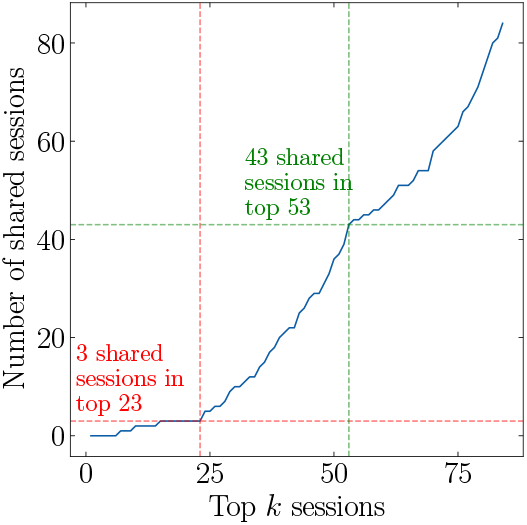
Number of shared sessions in the top-*k* ranking of all five heldout sessions.

## G Qualitative comparison on forward-prediction performances

Here, we qualitatively compare the forward-prediction results on some example neurons from Session 84. As shown in Figure 9, predictions made by the models trained on five topranked sessions (green) match the ground truth activity dynamics (blue) much better than those trained by five reverse-ranked sessions (orange). Neurons were randomly selected from the 100 most active ones.

**Figure 9:**
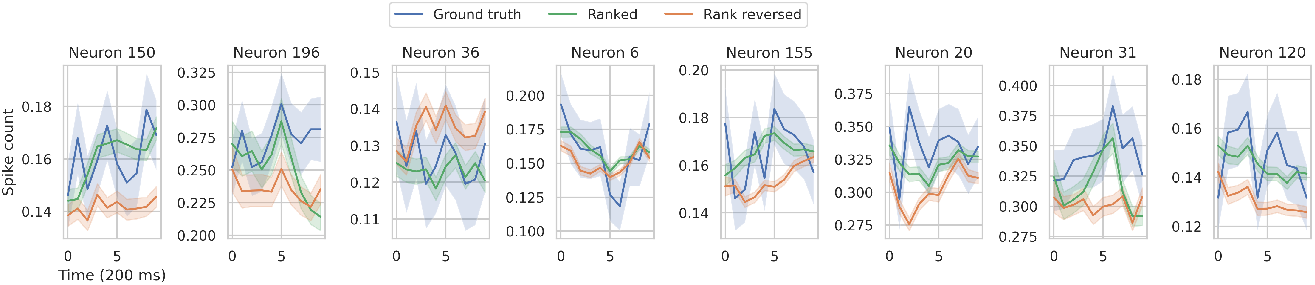
Qualitative comparison of forward-prediction performances between ranked and reverse-ranked models. Ground truth activities and predictions are averaged over trials. Shading shows standard error of the mean.

## Footnotes

1 Concurrent work on motor decoding by Ye et al. (2025) reports that the benefits from pretraining the model on 2000 hours of data are virtually nonexistent when finetuning datasets exceed 100 minutes, supporting our hypothesis that data heterogeneity issues extend beyond our dataset.

